# A Gradient of Hippocampal Inputs to the Medial Mesocortex

**DOI:** 10.1101/535047

**Authors:** Emanuel Ferreira-Fernandes, Carolina Quintino, Miguel Remondes

**Affiliations:** Instituto de Medicina Molecular, Faculdade de Medicina, Universidade de Lisboa, Lisbon, 1649-028, Portugal

**Keywords:** Hippocampus, anterior cingulate cortex (CG), retrosplenial cortex (RSC), medial mesocortex (MMC), hippocampal inputs, optogenetics, *in vitro* and awake-behaving multi-electrode recordings, long-range inhibitory neurons

## Abstract

Memory-guided decisions depend on complex, finely tuned interactions between hippocampus and medial mesocortical regions anterior cingulate and retrosplenial cortices. The functional circuitry underlying these interactions is unclear. Using viral anatomical tracing, *in vitro* and *in vivo* electrophysiology, and optogenetics, we show that such circuitry is characterized by a functional-anatomical gradient. While CG receives excitatory projections from dorsal-intermediate CA1 originated exclusively in *stratum pyramidale*, retrosplenial cortex also receives inputs originating in *stratum radiatum* and *lacunosum-moleculare*, including GAD+ neurons providing long-range GABAergic projections. Such hippocampal projections establish *bona fide* synapses throughout cortical layers, with retrosplenial cortex densely targeted on its layer 3, around which it receives a combination of inhibitory and excitatory synapses. This gradient is reflected in the pattern of spontaneous oscillatory synchronicity found in the awake-behaving animal, compatible with the known functional similarity of hippocampus with retrosplenial cortex, which contrasts with the encoding of actions and “task-space” by cingulate cortex.

**Highlights:** Both MMC regions CG and RSC receive monosynaptic connections from the dorsal-intermediate CA1

CG receives layer-sparse excitatory projections exclusively originated from *stratum piramidale* whereas RSC is targeted densely in superficial layers by a mixed excitatory and inhibitory input originating from all CA1 *strata*

CA1 monosynaptic projections correspond to active synapses onto distinct layers of the two MMC regions

Diverse synchrony between MMC and HIPP recorded *in vivo* is consistent with the rostro-caudal diversity of direct HIPP-MMC connections

## Introduction

Memory-guided decisions are characterized by complex, finely tuned interactions between the hippocampus (HIPP), and medial mesocortical (MMC) regions, namely anterior cingulate (CG), and retrosplenial (RSC) cortices (Alexander and Nitz, 2015; Battaglia et al., 2011; Cowen et al., 2012; Jones and Wilson, 2005; Rajasethupathy et al., 2015; Remondes and Wilson, 2013, 2015). Complex cognitive correlates of MMC neurons suggest an equally complex role of associating actions to outcomes in their episodic context, signaling errors and suppressing inappropriate responses (Carter et al., 1998; Shenhav et al., 2013; Vogt et al., 1992). Rodent lesion studies support a functional dichotomy between CG and RSC, with the latter absolutely critical for spatial memory-referenced behaviors (Aggleton, 2010; Hindley et al., 2014; Katche et al., 2013; Vann and Aggleton, 2005), and the former necessary for flexible choice behaviors and long-term memory (Chudasama et al., 2003; Goshen et al., 2011; Maviel et al., 2004; Teixeira et al., 2006; Vetere et al., 2011; Wang et al., 2012). These findings agree with the few studies recording MMC neural activity *in vivo*, and with the patterns of HIPP-MMC functional coordination (Remondes and Wilson, 2013, 2015). Neurons in CG encode choice effort (Cowen et al., 2012; Endepols et al., 2010) and task-space by assimilating into motor decisions the spatial-contextual properties of available choices (Remondes and Wilson, 2013; Sul et al., 2010), while RSC joins visual-spatial information such as head-direction and movement in space (Chen et al., 1994a, 1994b; Cho and Sharp, 2001) to encode spatial landmarks (Alexander and Nitz, 2015; Auger et al., 2012; Mao et al., 2017). The above findings are supported by the patterns of functional interaction between MMC and HIPP *in vivo*, namely the dynamic entrainment of RSC, CG and mPFC neurons by HIPP oscillations in the theta (Remondes and Wilson, 2013; Young and McNaughton, 2009), and gamma bands (Remondes and Wilson, 2015), both hypothesized to support the processing of relevant contextual information (Colgin et al., 2009; Girardeau et al., 2009; Jadhav et al., 2016; Pezzulo et al., 2014; Remondes and Wilson, 2013, 2015; Schomburg et al., 2014; Young and McNaughton, 2009). Hypothetically, diverse interactions of HIPP with distinct regions of MMC are supported by correspondingly diverse anatomical connections. These connections have been studied using distinct anatomical tracing molecules, and yielded results hard to reconcile, something critical to interrogate their role in neural synchrony and cognition. Some studies suggest the absence of hippocampal inputs onto CG (Jay and Witter, 1991), others its presence (Cenquizca and Swanson, 2007; Hoover and Vertes, 2007), exclusively from ventral HIPP (Cenquizca and Swanson, 2007), or show dorsal CA1 (dCA1), dorsal subiculum (dSUB) or CA1-SUB border monosynaptic projections to RSC (Cenquizca and Swanson, 2007; van Groen and Wyss, 1990, 1992; Van Groen and Wyss, 2003; Wyass and Van Groen, 1992). Furthermore, the physiological, or behavioral, relevance of such anatomical projections, namely whether they form actual *bona fide* synapses in CG or RSC that could be specifically manipulated, has never been shown. In sum, HIPP neurons with the potential of directly conveying contextual and mnemonic information onto the MMC, lending support to HIPP-MMC interactions found *in vivo*, are yet to be found. Here we decided to use anatomical tracing methods, *in vitro* electrophysiology combined with optogenetics, as well as *in vivo* electrophysiology in awake-behaving rats, to systematically study the global pattern of HIPP-MMC connectivity, identifying the neurons connecting the HIPP with the rostro-caudal divisions of MMC, and study the dynamic properties of such synaptic inputs.

Using Cholera Toxin subunit B conjugated with Alexa 647 (CTB-Alexa 647) injected in CG, mid-cingulate cortex (MCC) or RSC, we have found a population of labelled pyramidal neurons in dorsal-intermediate CA1 (diCA1) *stratum pyramidale* (SP) targeting CG, and a distinct population targeting RSC, distributed across pyramidal and sub-pyramidal CA1 *stratum radiatum* (SR) *and lacunosum-moleculare* (SLM), these last comprising GAD+ cells morphologically classified as interneurons. The HIPP inputs onto the region interfacing CG and RSC, globally classified as the MCC, intermediate an anatomical-functional gradient characterizing HIPP-MMC inputs. Furthermore, anterograde virus-mediated expression of hChR2.mCherry in diCA1 fills CG and RSC with distinctly distributed *labeled axons with terminal boutons* (LATB, resembling “beads-on-a-string”) (Cenquizca and Swanson, 2007), forming a rostro-caudal gradient of increasing density. Stimulating these terminals specifically with blue light while recording synaptic responses across MMC using a multi-electrode array (MEA, Multi-Channel Systems) allowed us to show that such HIPP terminals form region-specific *bona fide* synaptic inputs. Using sequential pharmacology and linear subtraction of responses (Chevaleyre and Siegelbaum, 2010) allowed us to isolate significant synaptic responses on all the divisions of MMC and to find that such responses are sensitive to the use of selective AMPA, NMDA and GABAa channel blockers. With this approach we isolated hippocampal long-range inhibitory synapses in RSC. Lastly, *in vivo* recording of multi-unit spikes simultaneously from HIPP and all divisions of MMC, revealed a gradient of HIPP-triggered spiking responses and synchronicity increasing rostro-caudally, consistent with the *in vitro* functional-anatomical data.

Thus, in summary, we found that hippocampal neurons establish functional synapses with all MMC divisions, wherein each rostro-caudal level of MMC receives diverse input from the full extent of diCA1. The fact that, contrary to CG, RSC receives a combination of excitatory and inhibitory long-range inputs from all CA1 *strata* is consistent with its functional proximity with HIPP, suggesting that RSC and HIPP jointly encode and store spatial context, while CG receives a ready-to-use spatial map to compute task-space in the service of decision-making.

## Results

### Contrary to CG, RSC receives monosynaptic projections from all diCA1 *strata*

We counted neurons retrogradely labelled by CTB-Alexa 647 following small-volume injections at individual rostro-caudal MMC coordinates (Figure 1A-B), and found diverse populations of labeled CA1 neurons, with each injection spot retrogradely labelling neurons along the full extent of diCA1 (Figure 1C-D), contrary to earlier reports (Cenquizca and Swanson, 2007; Jay and Witter, 1991). Most CA1 neurons targeting MMC originate in its dorsal-intermediate division (Figure 1E, ANOVA, *F*_(2,21)_=14.87, *p*=0.0001, n=10, followed by Bonferroni-corrected *post hoc* multiple comparisons), with no topography relating temporo-septal CA1 with rostro-caudal MMC (n-way ANOVA, *F*_(2,21)_=1.2, *p*=0.34, no significant interaction between MMC injection spot and number of neurons counted in each HIPP region). We then related the numbers of labelled neurons on each HIPP *stratum* (SO, SP, SR and SLM) with each cortical injection level (available in Supplementary Table 1), and found a significant difference in the HIPP *strata* providing inputs to the different levels of MMC (Figure 1F, n-way ANOVA, *F*_(3,28)_=11.94, *p*=0.000), as well as a significant interaction effect between the MMC injection level and neuron counts on each HIPP *stratum*, indicating that distinct rostro-caudal MMC regions receive inputs from distinct populations of HIPP neurons (Figure 1C-D and F, n-way ANOVA, *F*_(6,28)_=3.62, *p*=0.0087, there is a significant MMC injection level by HIPP *stratum* interaction). Specifically, injections of CTB-Alexa on CG labelled HIPP neurons located almost exclusively in CA1 SP (Figure 1C1and F, red boxplots, n-way ANOVA, all corrected p-values from multiple comparisons<0.0005). This result was confirmed by the injection of a non-selective retrograde virus rAAV2-retro-tdTomato (Tervo et al., 2016) in CG, which labeled a dense population of exclusively pyramidal neurons in all medial-lateral divisions of diCA1 (not shown). In contrast, RSC injections labelled pyramidal and non-pyramidal neurons across most diCA1 *strata* (Figure 1D1 and F, blue boxes, n-way ANOVA, all p-values from multiple comparisons ≅1, this result was also confirmed with a viral injection as above). The distribution of CA1 neurons labelled after CTB-Alexa injection in MCC resembles that seen for RSC, with no significant differences in the neuron numbers between *strata*, implying that MCC and RSC are indistinct in terms of HIPP connectivity (Figure 1F, green and blue boxes, n-way ANOVA, all p-values from multiple comparisons ≅1). However, the number of neurons labelled in SP after MCC injection is not different from the one obtained after CG injection (*p*=0.12), indicating an aspect of similarity between MMC and CG.

**Figure 1.**
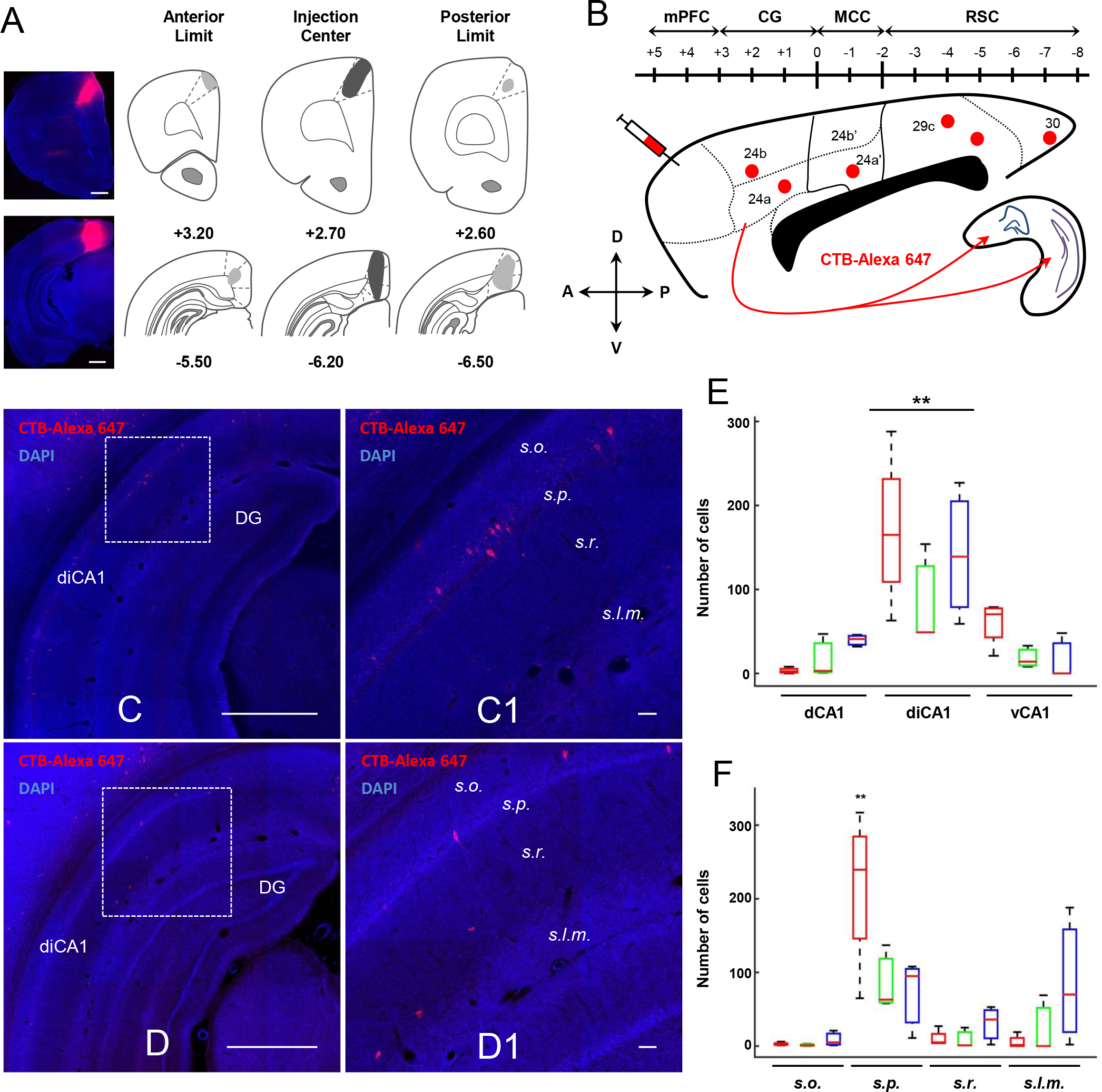
CA1 neurons directly project to the three main MMC regions CG, MCC and RSC. (A) Diagrams of center (black), anterior and posterior CTB-Alexa 647 injection extent (grey), in CG (top) and RSC (bottom). Magnification: 10x. Scale bars: 1000 μm. (B) Injection locations (red) across an MMC map. (C, C1) Retrogradely labeled hippocampal neurons in a representative slice of diHIPP, after the CG injection depicted in panel A. Higher mag in C1. Note CTB positive cells (red) restricted to CA1 SP (right). (D, D1) Retrogradely labeled hippocampal neurons in a representative slice of diHIPP, after the RSC injection depicted in panel A. Higher mag in D1. Note CTB positive cells (red) in SP, SR, and at the SR-SLM border (right). Magnification: 10x (C and D) and 20x (C1 and D1). Scale bars: 1mm (C and D) and 50 μm (C1 and D1). (E and F) Median and IQR of retrogradely labeled cells following injections in the CG (red, n=4 rats), MCC (green, n=3 rats), and RSC (blue, n=3 rats). On panel E cells were quantified according to their CA1 distribution in dorsal (dCA1), dorsal-intermediate (diCA1), and ventral (vCA1) HIPP, and on panel F according to hippocampal *strata* (SO, SP, SR, SLM). Asterisks indicate statistically significant differences (*p* < 0.05; n-way ANOVA). Note the presence of CA1 neurons directly projecting to all MMC regions with significantly higher numbers originating in diCA1. Note also that injections in CG label mostly neurons in SP whereas those in RSC label neurons throughout *strata*, and MCC shows an intermediate distribution. mPFC, medial prefrontal cortex. CG, anterior cingulate cortex; MCC, midcingulate cortex; RSC, retrosplenial cortex. See also Supplementary Table 1.

The above results demonstrate that there is an anatomical gradient in the source of hippocampal inputs targeting MMC, with RSC receiving inputs from all HIPP strata, and CG receiving inputs originating exclusively in CA1 pyramidal neurons. A distinctive feature of the above connectivity is its sparseness, discernible due to the small amounts of tracer injected, indicating that each restricted volume of MMC, corresponding to a limited population of neurons, receives information from the full dorso-medial hippocampal extent, providing MMC neurons with distinct epochs of a spatial representation by virtue of a phase-dependent representation of spatial trajectories (Dragoi and Buzsáki, 2006; Wilson et al., 2015), travelling medial-laterally across the longitudinal HIPP axis with a defined period (Lubenov and Siapas, 2009).

### Monosynaptic HIPP inputs to RSC include long-range inhibitory projections from GAD^+^ neurons at the SR-SLM border

Contrary to hippocampal projections to CG, those targeting RSC originate in neurons located in SP, but also interneuron-shaped neurons located in SR, SLM and at the SR-SLM border (Figure 3A), known to harbor a rich, behaviorally-relevant population of GABAergic neurons (Jinno et al., 2007; Lovett-Barron et al., 2014). If these are indeed GABAergic neurons, HIPP-RSC connections would include long-range inhibitory projections (LRIP) shown previously to originate in CA1-SUB to target RSC (Miyashita and Rockland, 2007), and also the entorhinal cortex, where they play a critical role in the processing of primary sensory inputs by the HIPP (Basu et al., 2016). To ascertain whether these are indeed GABAergic neurons, we performed GAD staining on the samples resulting from both CG and RSC injections. While we found GAD+ neurons widespread in all HIPP strata following injections on all regions, only following CTB-Alexa injections in RSC did we find neurons labeled with both CTB-Alexa and GAD (Figure 3A), and only at the SR-SLM border, (50 double-labelled neurons out of 90 retrogradely labelled with CTB-Alexa, in 6 analyzed slices from 3 animals) indicating that ~50% of all HIPP neurons projecting to the RSC, but not to CG, are indeed LRIP located at the SR-SLM border. These numbers agree with the ones resulting from recent findings (Yamawaki et al., 2018). The identification of LRIP between HIPP, EC and RSC suggest that this type of projection constitutes a trans-regional network, gating the transfer of high-level sensory information possibly involved in the storage of complex contextual information.

### Monosynaptic HIPP inputs differently target distinct levels of MMC

The previous retrograde labelling studies demonstrate the presence of distinct CA1 neuron populations targeting different levels of MMC but tell us nothing about the distribution of such projections at the destination, something crucial to understand the effect of such inputs on the cortical neural circuit. To investigate this, we injected AAV9.CaMKIIa.ChR2.mCherry in diCA1 to express the fluorophore mCherry in excitatory neurons (Figure 2A), and studied the anterograde expression of the fluorescent reporter at different MMC divisions (Figure 2B). While we found that this procedure resulted in the presence of noticeable fluorescence among layers L1-5, a conspicuous fluorescence peak centered in L3 was present in RSC, not pronounced in other cortical regions (Figure 2E and F, blue trace, compare to green and red plots). Furthermore, upon closer look, we found that mCherry fluorescence across MMC is due to the presence of sparsely distributed LATB (Cenquizca and Swanson, 2007) (Figure 2C, LATB, resembling “beads-on-a-string”) abundant in RSC, and gradually less dense as we move rostrally towards CG (Figure 2E). To quantify this observation, we manually counted the number of identified LATB on superficial (L1-4) and deep (L5-6) layers of each MMC region (see Methods) and compared such numbers, normalized by layer thickness. This quantification confirmed the presence of a gradient of HIPP-MMC connections, with a significant MMC region-by-layer (deep vs superficial) interaction (Figure 2D, n-way ANOVA, *F*_(2,1185)_=107.58, n=3 rats, *p*=0.0000), and a higher density of HIPP projections in RSC in its superficial layers, compared to deeper ones (*p*=0.0000). Following multiple comparisons, we found a significant increase in the number of LATB in the superficial, but not deep (*p*=0.12), layers of MMC as we move rostro-caudally (all *p*-values from multiple comparisons <0.01). These findings reinforce the notion of a rostro-caudal gradient of HIPP terminals across MMC regions.

**Figure 2.**
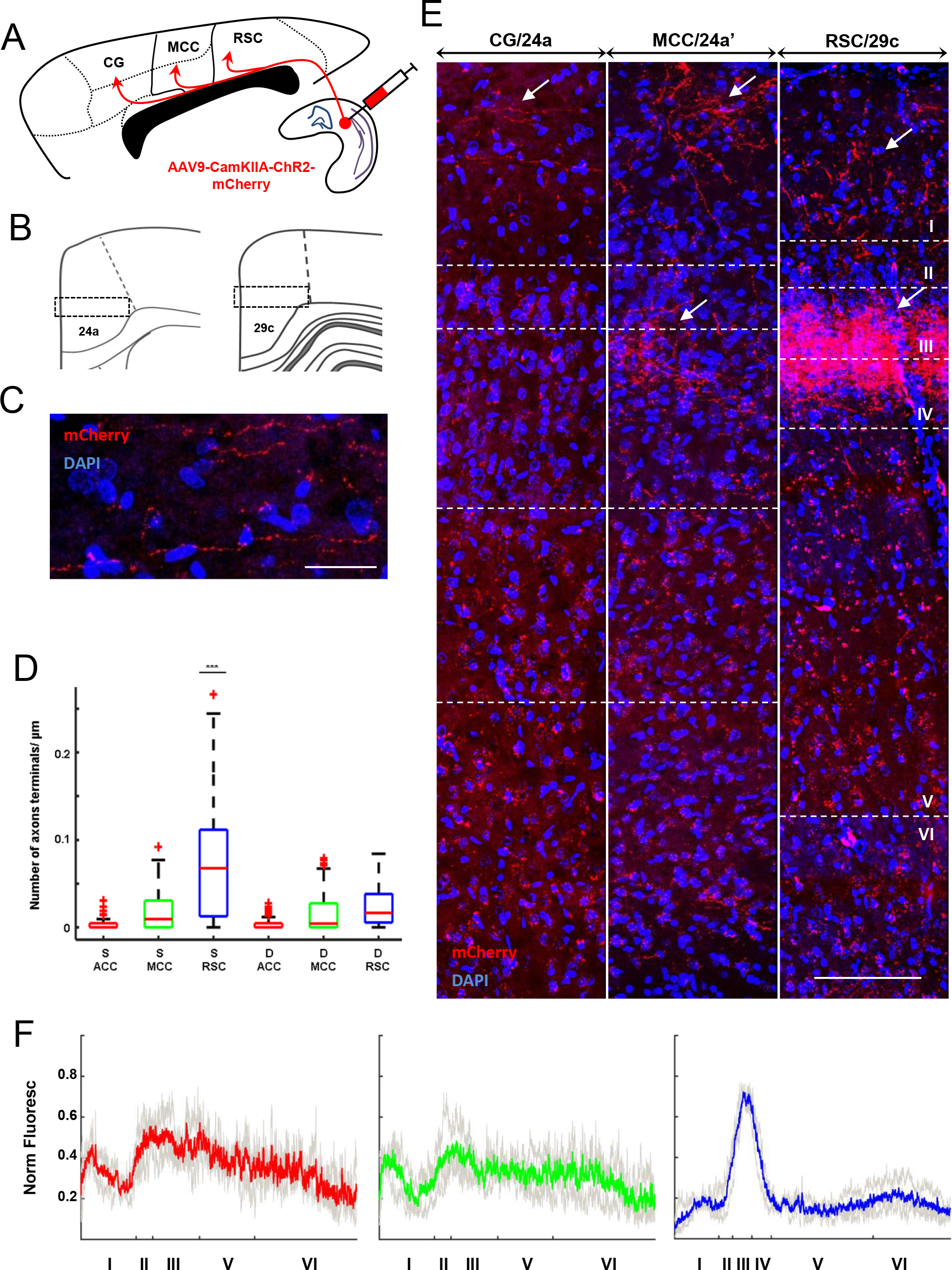
Laminar distribution of hippocampal excitatory inputs reveals a density gradient across the divisions of MMC. (A) AAV9-CaMKIIa-ChR2-mCherry injection locations in diCA1 for anterograde axon labelling. (B) Standard ROI used to quantify fluorescence distributions in MMC. (C) Confocal images of hippocampal excitatory terminals in MMC, taken from one illustrative rat brain expressing mCherry in diCA1. Magnification: 40x. Scale bar: 100 μm. Note the typical morphology of the sparsely distributed labeled axons with terminal boutons (Cenquizca and Swanson, 2007) (LATB, resembling “beads-on-a-string”). (D) Quantification of LATB on superficial vs deep layers of distinct levels of MMC showing increasing density towards RSC (*, *p*<0.01). (E) Illustrative ROI from the three MMC levels where LATB were counted, from CG (left) to RSC (right). White arrows point at mCherry+ LATB. (F) Normalized fluorescence across layers of CG (red), MCC (green), and RSC (blue) (n = 3 rats; red, green, blue, means; light gray, replicas).

**Figure 3.**
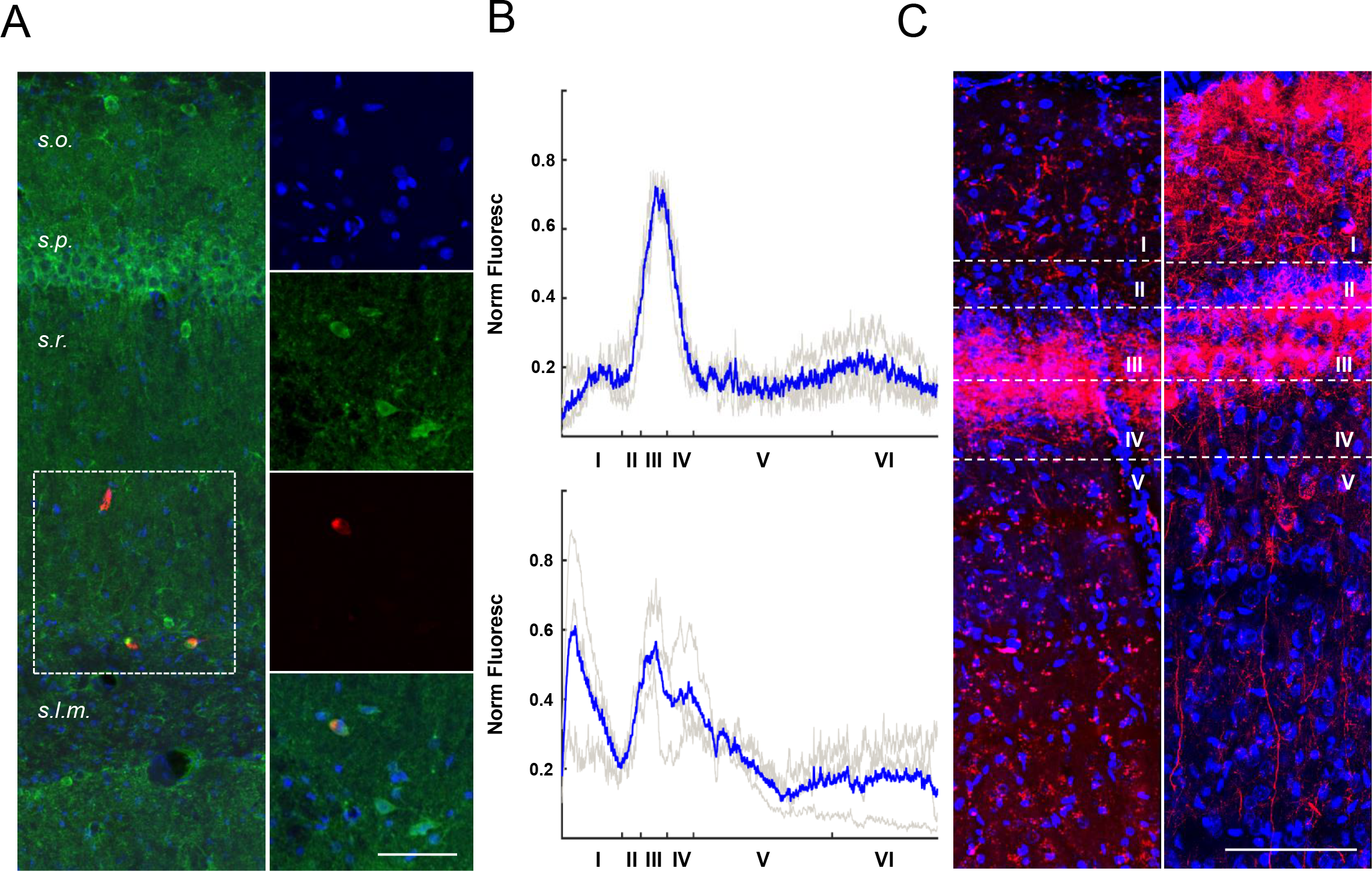
Retrogradely-labelled neurons in the CA1 SR-SLM border are LRIP projecting to layer 1 of RSC. (A) Injection of CTB-Alexa 647 in RSC labels neurons at the SR-SLM border (in red, as in Figure 1A), a subset of which (~50%) is also GAD^+^ and restricted to the SR-SLM border (magnification: 20x, scale bar: 50 μm, examples inside the red dotted square, one illustrative double-positive neuron in the four vertically arranged sub-panels, magnification: 63x, scale bar: 50 μm). (B) Normalized mCherry fluorescence distribution driven by either CaMKII (top panel), or the pan-neuronal promoter CAG, reveals distinct laminar distribution profiles, whose difference is attributable to the presence of LRIP axons originating in the HIPP. Note the extra peak in layer L1, in the bottom panel, absent from the profiles in the top panel (n=3 rats; *blue*, mean; *light gray*, replicas). (C) Illustrative RSC ROI depicting hippocampal terminals targeting RSC, following CaMKII (left column) or CAG-driven mChR2 expression. Magnification: 40x. Scale bar: 100 μm. See also Supplementary Figure 1, and Supplementary Table 2 for colocalization of mCherry+ with GAD+.

### LRIP originating in the SR-SLM border send monosynaptic inhibitory input to RSC

Since we found CA1 GABAergic neurons sending projections to RSC, we sought to describe and quantify the anterogradely-labelled putatively inhibitory axons targeting this cortical region. Assuming that the majority of non-pyramidal neurons establishing long-range connections with RSC are GABAergic, we used a pan-neuronal neurotropic virus expressing a fluorescent reporter (AAV9.CAG.ChR2.mCherry) and studied the distribution of resulting projections at the level of RSC (Figure 3B-C). We then compared the distribution of thus-obtained fluorescence-labeled terminals with the one found following the infection with CaMKIIa-specific virus, to infer the distribution of LATB corresponding to inhibitory terminals (Figure 3B-C). We found that a non-selective neurotropic virus injected in HIPP results in dense labelling of the border between the L3-4, such as the one obtained following CaMKIIa-promoted mCherry expression (repeated in Fig 3B-C, for comparison), but also a distinct, dense labelling on L1 (Figure 3B, bottom and C right panel) and on L2-3, resembling a third peak not seen in the CaMKIIa-promoted expression (Figure 3B, bottom plot, and C, right panel). Furthermore we noted the presence of vertically-oriented fibers in the deep layers of RSC, exclusively following infection with this pan-neuronal virus (Figure 3C, right panel, L4-5). Assuming all we have labelled in addition to the anterograde CaMKIIa-virus labelling originates in LRIP projecting to RSC, we have confirmed that CA1 sends LRIP to RSC layer L1, as previously described (Miyashita and Rockland, 2007). To further test whether this distinct labeling corresponds to inhibitory terminals targeting RSC, and in spite of the known technical challenges of labelling inhibitory axon boutons using GAD staining in brain slices (Temido-Ferreira et al., 2018), we performed such technique on the brain slices resulting from the above viral injections. We then analyzed the co-distribution of GAD+ and mCherry positive *puncta* using the Colocalization Threshold plugin from the Fiji Image analysis program (Temido-Ferreira et al., 2018). GAD+ puncta were found to colocalize significantly with mCherry-labeled axons among the dense labeling on L1 (Supplementary Figure 1, Supplementary Table 2), with an average tM GAD coefficient close to 0.5, suggesting a high proportion of co-occurring mCherry and GAD signals. In agreement with this, Costes’ threshold was in the [0, 255] range and the correlation below Costes’ threshold was around zero (Supplementary Table 2). This confirmed the presence of hippocampal LRIP in RSC layer 1. When the same analysis was performed in layers 3/4 and 5, the tM GAD coefficient was around 0.1 for layers 3/4 (Table 5) and the Costes’ threshold assumed negative values for layer 5 (data not shown), suggesting poor or absent colocalization in these layers. While the previous analyses support the notion that indeed the additional fluorescence peak we see in Figure 3B-C can indeed correspond to GABAergic terminals, we can no longer consider as such the GAD+ puncta located in the deeper layers (even though the detection of inhibitory axon boutons using GAD staining of neurites is notoriously difficult, raising the possibility of a false-negative result).

### Monosynaptic HIPP inputs to MMC constitute *bona fide* functional synapses

The presence of LATB originated in CA1 neurons and terminating in MMC suggests, but does not demonstrate, the existence of synapses established by CA1 axons onto MMC neurons, neither does it show how might such distinct distribution of synaptic inputs contribute to differences in HIPP-MMC neural synchrony (Alexander and Nitz, 2015; Remondes and Wilson, 2013). Local projections to the mPFC, CG and RSC have been studied *in vitro*, in slice electrophysiology experiments using local electrical stimulation, obtaining results that, while they might *include* the local synaptic response to stimulation of CA1 and dSUB-originated terminals (Hedberg and Stanton, 1995; Hedberg et al., 1993), fail in dissociating them from the multiple inputs crossing the same stimulating electrode location on their path, and coming from other brain regions. In order to specifically study the CA1 inputs targeting the different levels of MMC, and to ascertain whether they establish HIPP synaptic inputs, we infected diCA1 with AAV9.CaMKIIa.hChR.mCherry (Figure 4A-C), and prepared acute MMC slices from these brains for *in vitro* electrophysiology using a 64-electrode recording grid (MEA, Multi Channel Systems). We then recorded neural activity from the full rostro-caudal extent of MMC in individual slices, in response to stimulation of ChR2-expressing excitatory axons with a blue-LED light delivery system (Plexbrite Module, Plexon). Using this methodology, not only could we compare the local responses to stimulation of CA1 inputs across rostro-caudal levels of MMC, but also get an estimate of the events taking place *in vivo* when such stimulation is performed. We found that stimulation of CA1 terminals using 100 ms light pulses (@30mW total LED power) resulted in significant responses on all rostro-caudal levels, and region-specific distribution of responses in deep vs superficial layers (Figure 4D, red trace “CT”, and E color plots). Such responses include the depolarization due to the opening hChR2 channels while the light is on, which we separated from “pure” synaptic responses using sequential pharmacology (Figure 4 B, CT is control, followed by the distinct treatments T_CAM_ 1-3). Sequential addition of receptor-specific channel blocker drugs, the GABAaR blocker Picrotoxin (PTX, in T1_CAM_), the AMPAR blocker CNQX (PTX+CNQX in T2_CAM_), and the NMDAR blocker APV (PTX+CNQX+APV in T3_CAM_), leaves what constitutes the “hChR2-only” response (Figure 4D, blue trace) which we could then linearly subtract from the response obtained in the CT condition to obtain the pharmacologically suppressed component, i.e. the synaptic response (Figure 4D, black trace). We could thus study its initial slope and amplitude, across the layers of distinct MMC levels (Figure 4E-F). We found significant synaptic responses (p<0.05; Wilcoxon rank sum test comparing distinct pharmacological conditions) in many of the MEA channels, and distinct patterns of activation depending on the MMC rostro-caudal level analyzed. While there are no significant global differences in the slope and amplitude in the deep vs superficial cortical layers, nor on CG vs RSC responses, we found a significant layer-by-region interaction (Figure 4F, Slope, n-way ANOVA, *F*_(1,38)_=10.31, *p*=0.0027 and Amplitude, *F*_(1,38)_=11.55, *p*=0.0016, n=8 rats), indicating that the slope and amplitude of the deep vs superficial responses to CA1 input stimulation depends on the MMC rostro-caudal level, with CG receiving stronger synapses in its deep layers, and the opposite occurring in RSC (Figure 4E). Analysis of absolute values largely abolished the differences observed, further confirming their dependence on layer-specific dipoles, reflecting, at least partially, the anatomical differences observed before. In fact, the patterns of responses closely match the density of LATB seen in Figure 3, with strong responses in the superficial layers of RSC, densely targeted by CA1 projections, and stronger responses in deeper layers of CG, which harbor sparse CA1 terminals.

**Figure 4.**
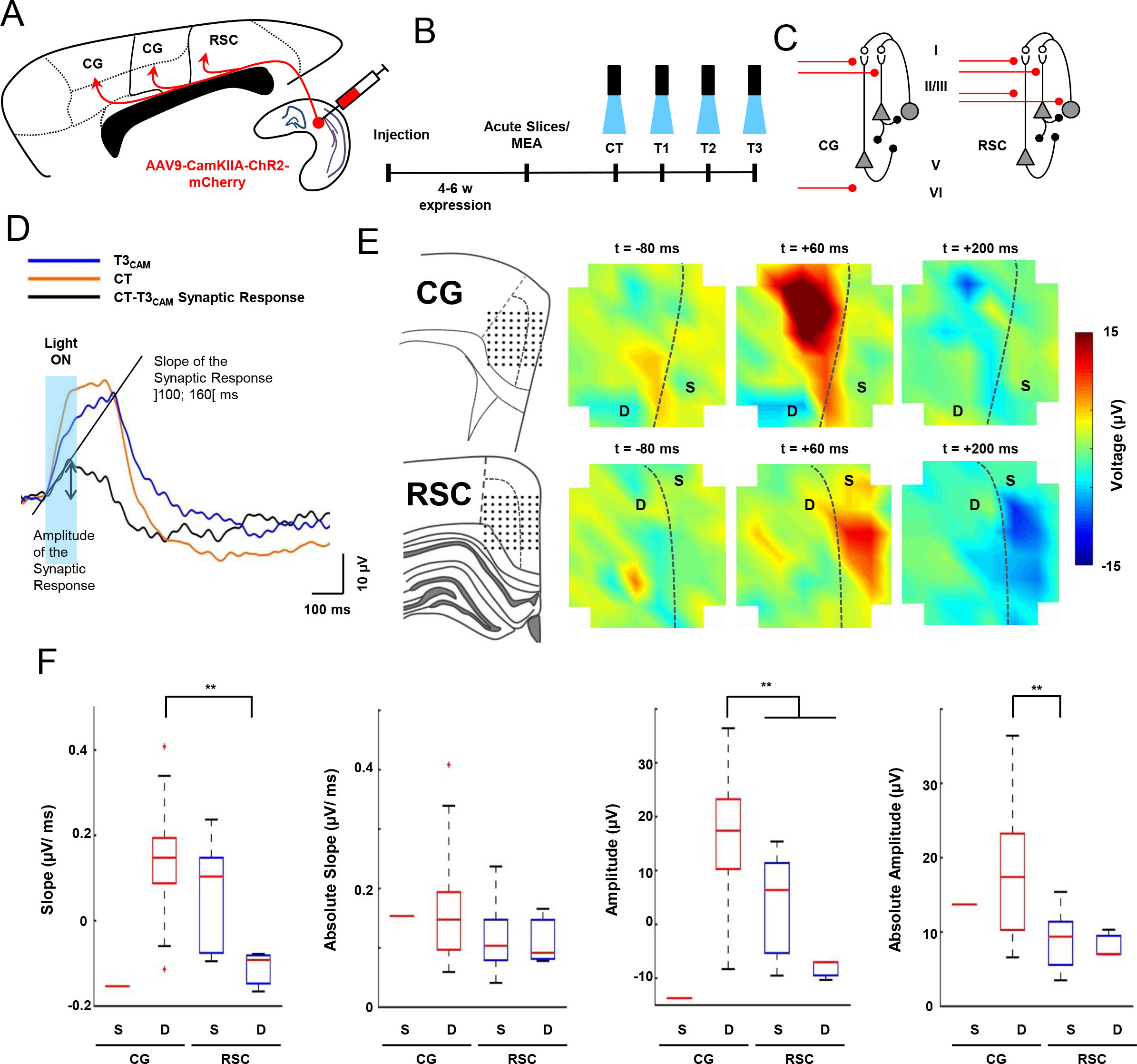
Direct hippocampal projections to the MMC establish *bona fide* hippocampal-cortical synapses. (A) Diagram of the experimental protocol wherein following the injection of AAV9.CaMKIIa.hChR2.mCherry in diHIPP, (B) acute slices were cut and placed on a MEA array to record synaptic responses to blue light optogenetic stimulation of local hippocampal terminals, during multi-site electrophysiological recordings, and sequential pharmacological treatments (see Methods, CTRL-CT, T1_CAM_-PTX, T2_CAM_-PTX+CNQX, T3_CAM_-PTX+CNQX+APV) (C) Scheme of the CG and RSC circuits to be stimulated, CA1 inputs labelled red. (D) Representative ChR2-evoked extracellular response explaining the computation of the synaptic component by linear subtraction of responses under distinct drug conditions (CT – T3). Each color-coded trace corresponds to the response obtained in one MEA contact (averaged over 2 min) in a CG slice, under control (CT) and T3_CAM_ treatment (T3). Computation of slope and amplitude of the synaptic response are illustrated (see Methods). (E) Diagrams depict the apposition of the MMC slice onto the MEA electrode array. Colorplots are representative MEA voltage amplitudes of the synaptic responses in CG (top) and in the RSC (bottom) at −80 ms, +60 ms and +200 ms of blue light onset. Note the distinct anatomical distribution of response amplitudes in CG vs RSC. The dotted line on all panels separates superficial and deep layers. (F) Boxplots of synaptic response slope, absolute slope, amplitude and absolute amplitude in CG and RSC (n = 4 rats for each brain region, * *p*<0.05). CG responses are more intense than RSC ones, and higher in deep layers, contrary to RSC.

The above findings are evidence that the topographical gradient of HIPP inputs translate into functional rostro-caudal differences in the MMC’s synaptic responses, lending strong support to the notion held previously that anatomically defining brain regions is more than simply a matter of morphological taxonomy, but rather constitutes modelling the brain, as it generates distinct predictions concerning its functions (Vogt, 2016).

### Long range projections from GAD+ HIPP neurons to RSC constitute functional GABAergic inputs

Having identified LRIP originating in CA1 and specifically targeting RSC, we sought to ascertain whether these neurons do establish functional GABAergic synapses. This is a critical first step to understand the functions of such neurons in the context of HIPP-MMC circuit, as well as its consequences for behavior. To manipulate inhibitory projections onto MMC we used the pan-neurotropic virus mentioned before (AAV9.CAG.hChR2.mCherry), and used a modified version of the pharmacological protocol used before to isolate such putative input. By placing PTX at the end of the sequence, we first blocked excitation using CNQX+APV (treatment T2_CAG_), hence blocking local feedforward inhibition and sparing only the non-local (CA1-originated) PTX-sensitive inhibitory component, plus the hChR2-resulting potential. We then added PTX, leaving only the hChR2 component, which we could subtract from condition T2_CAG_ to obtain a pure GABAergic response originated remotely (Figure 5A-C). These responses are depicted in the top panels of Figure 5D and E. Once the hChR2 plus inhibition component, obtained by T2_CAG_ is subtracted from the control trace, we obtain the distribution of excitatory input, essentially concentrated in the superficial layers, as found previously. T3_CAG_ (CNQX+APV+PTX) then fully blocks excitation plus fast inhibition (Chevaleyre and Siegelbaum, 2010), leaving only the optogenetic depolarization response (hChR2), and thus its subtraction from condition T2_CAG_ (optogenetic response plus non-local inhibition) will isolate the non-local GABAergic response, depicted in the lower panel of Figure 5D and in the lower plots of Figure 5E. We thus found that the MMC response triggered by the HIPP LRIP can indeed be isolated and studied, its slope and amplitude are both significantly lower than the ones corresponding to excitatory responses (n-way ANOVA, Slope, *F*_(1,54)_=66.9, *p*=0.0000 and Amplitude *F*_(1,54)_=31.47, *p*=0.0000), and they tend to distribute around the superficial layers. While the amplitude of the responses tends to favor electrode contacts classified as “deep” in the previous experiment (n-way ANOVA, Amplitude, *F*_(1,54)_=22.14, *p*=0.0000, Bonferroni corrected for multiple comparisons, excitatory amplitude, *p*=0.0031, inhibitory amplitude *p*=0.0033), such contacts are located close to the cortical surface, something not seen when using the CaMKIIa-driven hChR2 (Figure 5F). This is in agreement with previous anatomical findings (Miyashita and Rockland, 2007; Yamawaki et al., 2018), as well as our own, concerning the anatomical distribution of pan-neuronal terminals, in that there are indeed projections targeting the superficial RSC layers, but also conspicuous, vertically-oriented terminals (Figure 3C, right panels), possibly providing GABAergic input to both the superficial and the deep sides of layers L3-4. Finally, we noted that the PTX-sensitive conductance we now describe has the same polarity as the CNQX and APV sensitive ones (Figure 5D-E). Several explanations are possible for this. We can be systematically recording A) positive going potentials due to neuronal hyperpolarization, something hard to reconcile with positive going excitatory potentials measured locally, B) long-range inhibitory projections acting as a disinhibitory gate that transiently suppresses feedforward inhibition (Basu et al., 2016), or C) eliciting excitatory GABAergic currents (Gulledge and Stuart, 2003), something reported to occur in somatic and dendritic GABAergic responses under physiological conditions that preserve intracellular anion concentration (such as extracellular recordings), once such inputs are sufficiently isolated from excitatory inputs (Andersen et al., 1980; Stein et al., 2003). We cannot exclude either of the two last possibilities.

**Figure 5.**
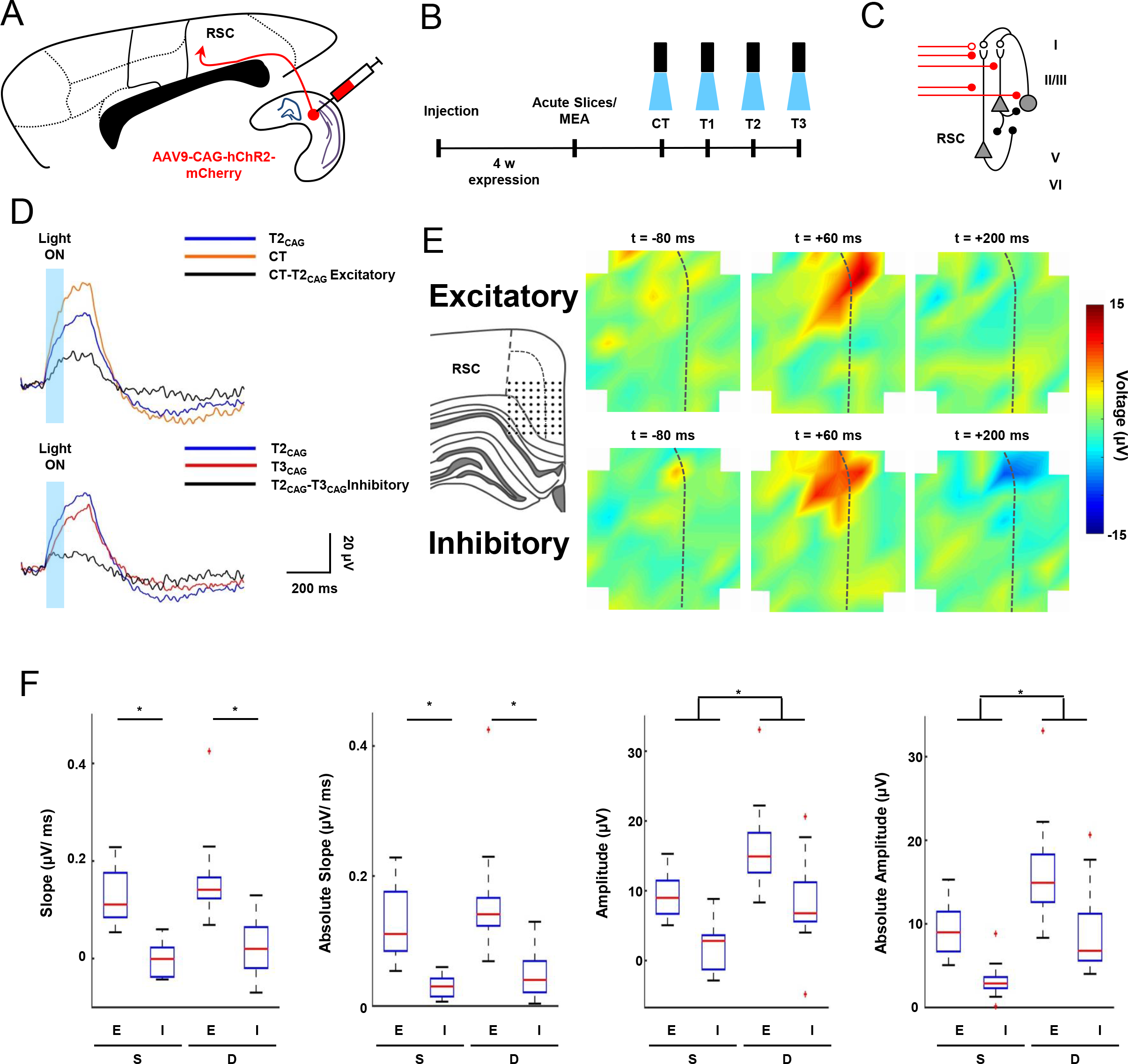
CA1 LRIP projecting to RSC establish *bona fide* hippocampal-cortical GABAergic synapses. (A) Diagram of the experimental protocol wherein following the injection of AAV9.CAG.hChR2.mCherry in diHIPP, (B) acute slices were used to record synaptic responses as above with sequential pharmacological treatments adapted (CTRL-CT, T1_CAG_-APV, T2_CAG_-APV+CNQX, T3_CAG_-APV+CNQX+PTX) to dissociate the excitatory (CT-T2_CAG_) and inhibitory (T2_CAG_-T3_CAG_) synaptic responses (C) Scheme of the RSC circuit to be stimulated, open circle represents inhibitory long-range inputs. (D) Representative example as above, adapted to isolate the excitatory (CT_CAG_-T1_CAG_) and inhibitory (T1_CAG_-T2_CAG_) synaptic responses. Each color-coded trace corresponds to responses in control (CT), T1_CAG_, and T2_CAG_ treatments. (E) Diagrams depicting an RSC slice apposed on the MEA electrode array. Colorplots depict synaptic responses in RSC at −80 ms, +60 ms, and +200 ms, for excitatory (top row) and inhibitory (bottom row) responses. (F) Boxplots of synaptic response slope, absolute slope, amplitude and absolute amplitude (n = 4 rats, *, *p*<0.05). Note the presence of a significant inhibitory response to the stimulation of hippocampal axons, whose absolute amplitude is higher in layers right below the superficial ones (bottom color plot in E).

### *In vivo* hippocampal-triggered MMC neural activity is consistent with the presence of diverse monosynaptic connectivity between HIPP and distinct levels of MMC

Having found that the HIPP directly connects to the distinct levels of MMC, establishing *bona fide* synapses therein, and that these connections support diverse remote-to-local neural circuitry, we sought to study the pattern of HIPP-MMC spontaneous co-activity that might result from such diversity. We thus recorded multi-unit neural spiking activity (MUA) from the diCA1, simultaneously with the full extent of MMC, encompassing CG, MCC and RSC, on awake rats in various behavioral conditions, and compared cortical responses to HIPP spikes. To compute such responses we extracted 2 sec epochs of z-scored cortical MUA triggered by 10 ms time bins containing at least 4 HIPP MUA spikes. As a control, we compared these epochs with equivalent ones triggered by randomly sampled time bins. We found that cortical responses to HIPP spiking are significantly higher than randomly picked epochs, on all 3 MMC regions (Figure 6, from 12 datasets, *t*_(20,16,22)_=3.04, 3.41, 2.77, *p*=0.006, 0.004, 0.01, for CG, MCC and RSC respectively), with no significant differences between distinct regions. A significant increase in cortical activity within 10 ms of HIPP spiking is consistent with a long-range monosynaptic connection. To achieve finer temporal resolution, and even though the raw local field potential (LFP) is prone to volume conduction, as well as other artifacts, we wanted to see whether we could observe qualitatively noticeable changes in the LFP in response to HIPP spikes. We extracted raw LFP data segments triggered on the same time bins defined above, and indeed noticed complex deflections on all three regional responses (Supplementary Figure2). We couldn’t help noticing that, like in the MUA response, the amplitude of RSC LFP responses to HIPP spikes is somewhat smaller than the ones registered in more rostral MMC levels, CG and MCC. Such is suggestive of a more constrained RSC depolarization in response to HIPP spikes, possibly brought about by the co-activity of long-range inhibitory projections, in parallel with excitatory ones.

**Figure 6.**
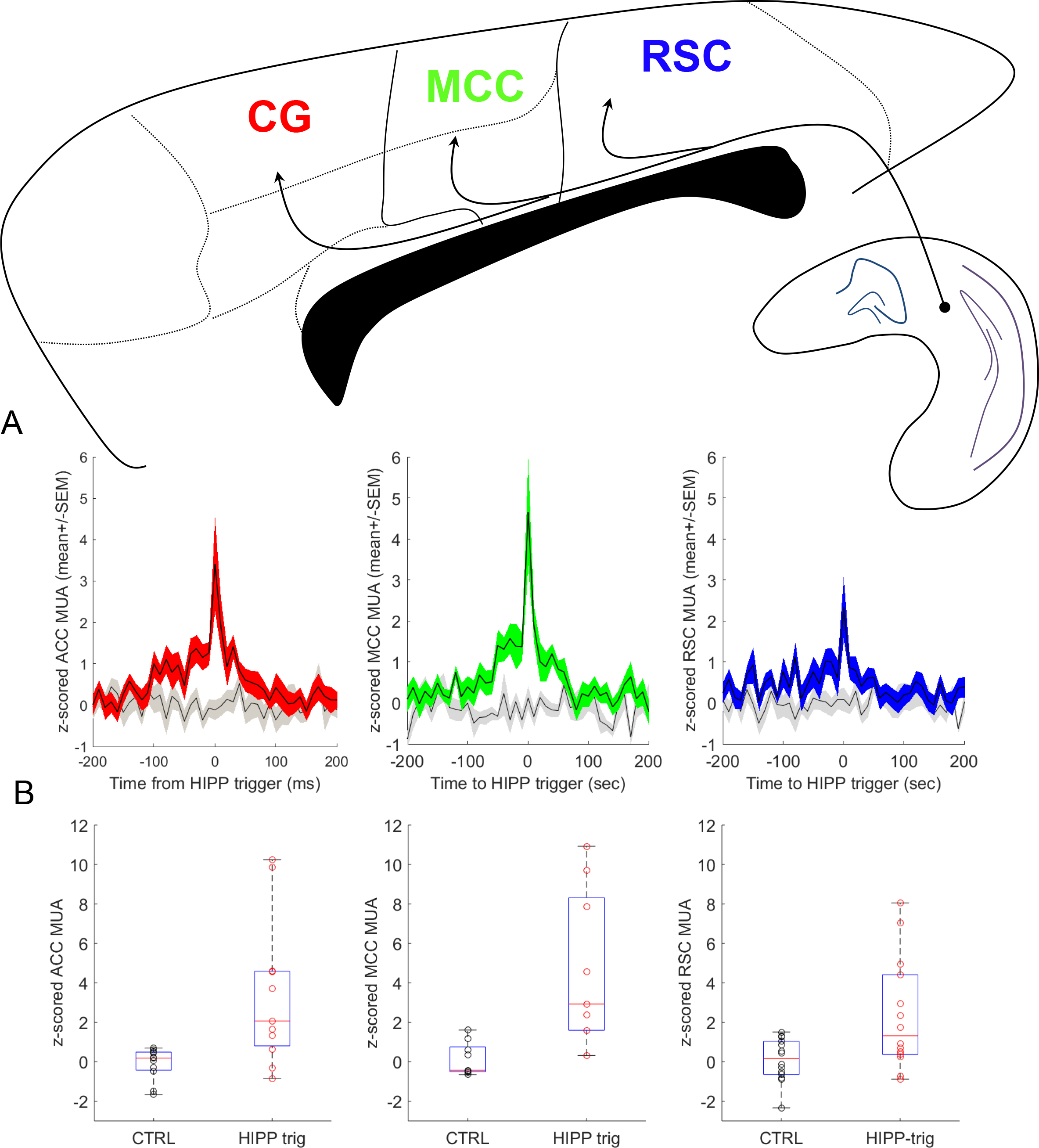
Hippocampal-triggered cortical activity *in vivo* is consistent with the presence of direct connectivity of CA1 with MMC regions. (A) Color-coded plots depict CA1-triggered MMC MUA average+/−SEM, overlaid onto grey-colored plots depicting cortical MUA triggered by randomly chosen time bins. Note the presence of increased activity on all MMC regions, within 20-30 ms of the trigger point, somewhat lower in RSC. (B) The row of boxplots depicts cortical MUA across datasets (median+/-IQR), and the randomly-triggered for comparison. All regions exhibited statistically significant MUA increases (*p*<=.01). See also Supplementary Figure 2.

The diverse connectivity we have reported above, and the very conspicuous MMC responses to HIPP spikes prompted us to ask whether such diversity could signify distinct patterns of activity, namely oscillations, and dominant modes of neural coordination. To answer this question, we performed a spectral analysis of the HIPP-triggered MUA binned at 10 ms like before, on a 500 ms sliding window stepped every 50 ms. A qualitative assessment of the three regions’ spectrograms, depicted in Figure 7 shows the presence of power increases on the main biologically-relevant frequencies reported in hippocampal-cortical ensembles, namely theta (5-8 Hz), beta (13-18Hz), slow-gamma (23-31 Hz) and fast-gamma (40-50Hz), something absent from the control data (in Figure 7 shown only for hippocampal MUA). Furthermore, there seems to be a rostro-caudal gradient in the dominant response frequencies of population neural activity, with increasing relative power at progressively higher frequencies as we move caudally in cortex (Figure 7B, note distinct color scales). This gradient of spectral power is accompanied by distinct patterns of oscillatory synchrony, as measured by the spike-triggered HIPP-MMC coherence of the binned MUA (Figure 7C, distinct color scales). To quantify and test this hypothesis, we normalized power and coherence to a pre-trigger baseline of 0.5 seconds, took the mean at each of the above frequency bands on each dataset, (Figure 8 A-B and S3), and compared their magnitude in the half-second before the trigger vs the half-second after the trigger, across MMC using n-way ANOVA (Figure 8B). This analysis revealed a HIPP-triggered power increase in all regions of MMC, regardless of MUA frequency (Figure S3B, *F*_(1,224)_=78.06, *p*=0.0000, n=11 datasets, we present here boxplots for each condition for informative purposes). We then sought to investigate whether there were regional differences in the relative magnitude of coherence across the frequencies analyzed. For this, we used the same normalization procedure as above, and compared magnitudes at the same time points and frequencies as above (Figure 8A). We found a main effect of the HIPP trigger (Figure 8B, *F*_*(2,224)*_=26.37,*p*=0.0000, n=11 datasets, we present here boxplots for each condition for informative purposes) confirming the presence of a HIPP-triggered increase in HIPP-MMC synchrony. In addition, we found a significant trigger vs region interaction (*F(2,224)*=3.92, *p*=0.021), indicating that such synchrony depends of the cortical region analyzed. Post-hoc comparisons revealed a region-related gradual increase in the mean coherence, culminating with a significant HIPP-triggered increase in coherence between HIPP and the posterior most RSC (*p*=0.0000, Bonferroni), itself significantly different from the post-trigger coherence between HIPP and CG (*p*=0.02, Bonferroni).

**Figure 7.**
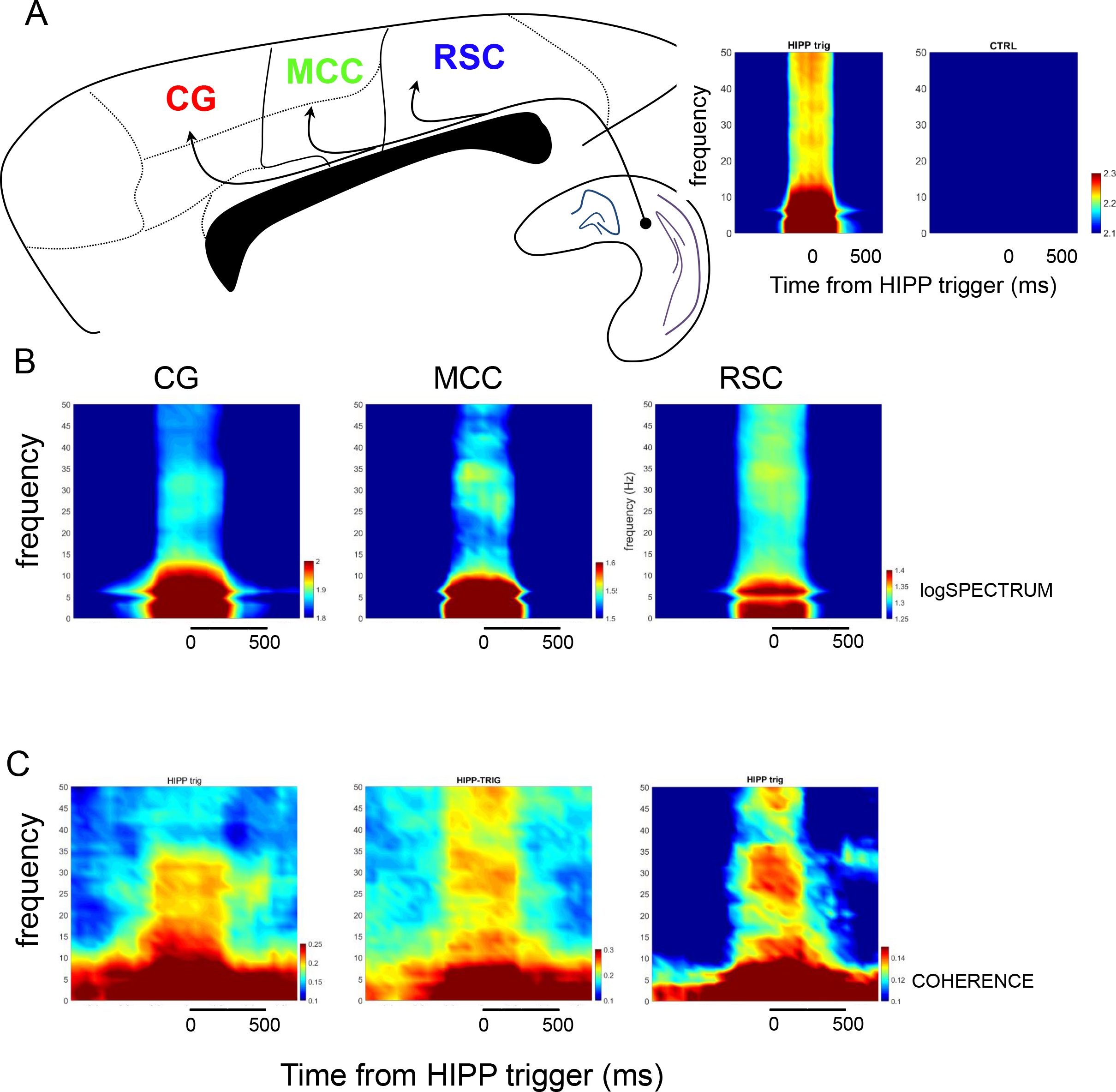
Hippocampal-cortical spectrograms and coherograms support the presence of short-latency oscillatory synchrony. (A) Diagram of the MMC regions and HIPP, and (right) spectrogram of the CA1-triggered MUA. Note the presence of increased power at all relevant frequencies, and the absence thereof in the control data. (B) HIPP-triggered MMC MUA spectrograms. Note the presence of increased power at behavior-relevant frequencies in all regions. (C) CA1-triggered MMC MUA coherograms illustrate the temporal alignment of CA1 and MMC MUA. Note the presence of coherence in all MMC regions, somewhat clearer in RSC.

**Figure 8.**
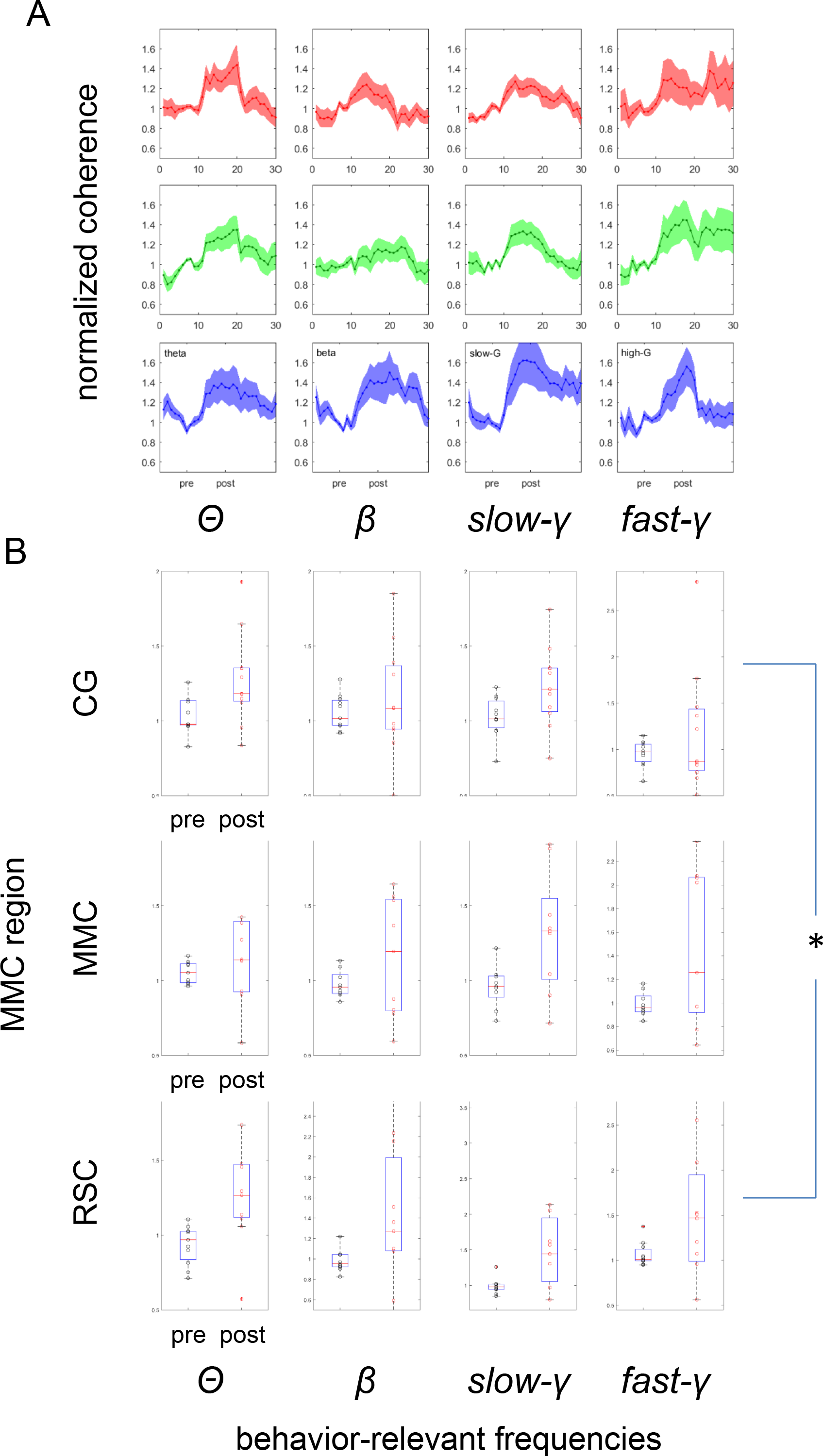
HIPP-triggered HIPP-MMC coherence increases specifically in the RSC. (A) Color-coded plots depict the quantification of CA1-triggered CA1-MMC coherence at relevant frequency bands, normalized to a pre-trigger baseline of 0.5 seconds. (B) CA1-triggered CA1-MMC coherence significantly increases across MMC (*p*=0.0000, n=11 datasets, we present here boxplots for each condition for informative purposes), with a significant trigger vs region interaction, and a region-related gradual increase in the mean coherence, culminating with a significant HIPP-triggered increase in coherence between HIPP and RSC (*post-hoc* comparisons, *p*=0.0000, Bonferroni), itself significantly different from the post-trigger coherence between HIPP and CG (*, *p*=0.02). This anatomically-distributed coherence pattern is suggestive of a rostro-caudal gradient underlying the communication between the HIPP and the distinct divisions of MMC. See also Supplementary Figure 3.

## Discussion

A major obstacle in dissecting the neural bases of memory-guided behavior lies in reconciling multiple anatomical studies describing HIPP-MMC interconnections, something critical for their interrogation in the context of cognitive mechanisms. We have used retrograde and anterograde anatomical tracing techniques to compare the connections between HIPP and different anatomical levels of the MMC, CG through RSC. This systematic study provides anatomical support to distinct functional proximity between either CG and RSC, and the HIPP. While the full extent of MMC receives monosynaptic inputs from the diCA1, each region of MMC receives distinct HIPP inputs with diverse layer distributions. RSC receives a stronger HIPP input originating from all HIPP *strata*, something reflected in their functional inter-dependence and coding properties, wherein RSC neurons exhibit significant activity changes in response to HIPP lesions (Albasser et al., 2007), and respond to visual-spatial variables necessary for contextual memory processing, such as HIPP-like place-selectivity (Mao et al., 2017). RSC would thus access a “raw” version of the HIPP output, whose information both jointly process at an early stage of spatial mapping. Conversely, CG receives HIPP inputs originating exclusively in CA1 pyramidal neurons, thus reading an already processed spatial map, in the service of task-space coding and behavioral control, consistently with previous *in vivo* studies (Remondes and Wilson, 2013, 2015; Yu and Frank, 2015). The distinct anatomical connectivity of CG and RSC with HIPP is intermediated by the one with MCC which includes aspects of both. The dichotomy we now show is consistent with lesion studies showing that, contrary to CG, lesions in RSC critically affect performance in tasks that rely on contextual memory, (Maviel et al., 2004; Vann and Aggleton, 2002). It is notable that each restricted volume of MMC, corresponding to a limited population of neurons, “reads” information from the full dorsal-intermediate hippocampal extent, thus providing cortical neurons with distinct epochs of an episodic representation by virtue of a phase-dependent representation of place-sequences (Dragoi and Buzsáki, 2006; Wilson et al., 2015), travelling medial-laterally across the longitudinal HIPP axis with a defined period (Lubenov and Siapas, 2009). This implies that restricted groups of CG and RSC neurons have access to the spatial and temporal content of a global episode, encoded in HIPP.

Using multi-electrode *in vitro* electrophysiology combined with optogenetics and sequential pharmacology, we isolated significant synaptic responses on all the divisions of MMC in response to stimulation of HIPP terminals, and found that such responses are sensitive to the use of selective AMPA, NMDA and GABAa channel blockers, like *bona fide* synapses, and also that the distribution of thus-analyzed responses is consistent with the anatomical distribution of LATB present at each MMC level.

Our results bring new light to an old controversy. Previous studies suggest the absence of dorsal hippocampal inputs onto CG (Jay and Witter, 1991), others their presence (Cenquizca and Swanson, 2007; Hoover and Vertes, 2007), and also that most HIPP projections towards RSC reportedly originate in the contiguous dSUB or from neurons in the CA1-SUB border (Cenquizca and Swanson, 2007; van Groen and Wyss, 1990, 1992; Van Groen and Wyss, 2003; Wyass and Van Groen, 1992).

By systematically analyzing HIPP-MMC connectivity, we now present a quantitative account of hippocampal inputs to MMC divisions. We show that HIPP and MMC are indeed connected directly by a population of neurons from diCA1, following a caudo-rostral gradient in which a dense, dual (excitatory/inhibitory) and layer-specific projection is progressively converted in a sparse, excitatory, and diffuse projection. These observations suggest that hippocampal activity informs CG and RSC computations at different levels. RSC receives multi-layer hippocampal inputs mainly in its superficial layers (L1-3), where it sends and receives most corticocortical connections, and CG is targeted exclusively by CA1 pyramidal neurons with stronger potentials evoked in L5 whose large pyramids project descending axons to the striatum and other subcortical structures, consistent with executive functions and behavior control. By using *in vivo* multi-site recordings of both neuronal spikes and LFP, we have shown that the spontaneous activity patterns in the HIPP and MMC in the awake-behaving rat follow what would be expected from the above-described connectivity. First, epochs of increased spiking from HIPP, are accompanied by short-term increases in MMC areas, with increased levels generally preceding and following the trigger point, indicative of complex time-dependent cross-talk between these regions (Jadhav et al., 2016; Kay et al., 2016; Remondes and Wilson, 2015; Yu et al., 2017). Second, such increases are somewhat stronger in the anterior-most regions of MMC, something also apparent from LFP data analyzed in a similar manner, implying that the presence of LRIP in parallel with excitatory HIPP inputs modulates RSC cortical responses *in vivo*. Our *in vivo* data further shows that MMC responses to HIPP spikes have an oscillatory component favoring frequencies known to play a significant role in hippocampal-cortical functions (Bieri et al., 2014; Buzsáki and Moser, 2013; Colgin and Moser, 2010; Engel and Fries, 2010; O’Keefe, 1993; Zheng and Colgin, 2015). Contrary to the HIPP-triggered increase in cortical firing rate, the strength of oscillatory alignment to the HIPP rhythms increases gradually as we move caudally along the MMC divisions, with the posterior-most RSC regions significantly more engaged to the hippocampal oscillations, which is consistent with RSC receiving denser HIPP input, from all HIPP layers, both excitatory and inhibitory. This is especially relevant since inhibitory inputs have widely been considered the main effectors of gamma oscillations and long-range gamma synchrony (Chen et al., 2017; Jinno et al., 2007; Mann and Paulsen, 2007; Paulsen and Moser, 1998; Traub et al., 1996), namely during working memory-dependent behaviors (Abbas et al., 2018).

The fact that RSC receives long-range excitatory and inhibitory inputs from all HIPP *strata*, matched by enhanced HIPP-RSC synchrony, is consistent with the notion of continuous feedback and functional proximity between RSC and HIPP, jointly processing spatial landmarks in the service of contextual memory and spatial navigation. Conversely, more diffuse hippocampal inputs to GC, with stronger potentials evoked around layer 5, would result in stronger responses guiding downstream executive behaviors via large pyramidal neural projections to the basal ganglia.

## Experimental Procedures

### Animals

All procedures were performed in accordance with EU and Institutional guidelines. For anatomical, *in vitro*, and *in vivo* electrophysiology, we used 16, 11 and 2, respectively, male rats aged 3-6 mo.

### Retrograde and anterograde tracing

CTB-Alexa 647 (Molecular Probes) was diluted in sterile PBS (0.5% (wt/vol)) and micro-injected (500 nL) in each MMC spot (all coordinates listed in ST3). Rats were sacrificed and perfused 7-11 days after injection for further processing (Conte et al., 2009; Varela et al., 2014). Vectors AAV_9_.CaMKIIa.hChR2-mCherry, or AAV_9_.CAG.hChR2-mCherry were micro-injected in diHIPP, rats were sacrificed after 30 days (Morgenstern et al., 2016). All constructs were acquired under appropriately signed MTAs.

### Stereotaxic surgical injections

Rats were anesthetized and prepared for craniotomy as previously published (Remondes and Wilson, 2015). The skin was then incised and retracted to expose the skull where a craniotomy and durotomy were performed. The tip of a glass micro-pipette loaded with the tracer was lowered into the brain where the appropriate agent was injected at 50 nL/minute, left in place for 15 min, and removed at 0.5 mm/min. The wound was sutured, and the rat was allowed to recover (protocol based on Conte et al., 2009).

### Euthanasia, perfusion, histology, immunohistochemistry

After perfusion the brains were placed in 4% PFA for 24 hours at 4°C, equilibrated in 30% sucrose 4% PFA, embedded in gelatin, and frozen. Coronal sections (100 μm) were cut in a cryostat (Leica, CM3050 S). Sections were incubated in 1μg/ mL of DAPI (Sigma) for 20 min and mounted in Mowiol (Sigma) or (neurochemistry) 0.1 M PBS 0.5 % Triton X-100 for 15 min followed by 0.1 M PBS 10% rabbit serum (RS) 1% (BSA) for 1h. Sections were then incubated in anti-GAD65/67 (Sigma-Aldrich, G5163), washed 3×15 min in PBS, and placed in goat anti-rabbit IgG-Alexa Fluor 488 (ThermoFisher, A-11008). Stained sections were washed 3×15 min in PBS, incubated in 1μg/ mL of DAPI for 20 min and mounted in Mowiol (Miyashita and Rockland, 2007).

### Microscopy and Image analysis

Coronal slices were imaged on an Axio Observer microscope (Zeiss) coupled to an Axiocam 506 mono CCD (Zeiss). Zeiss filters 49 and 50 were used to observe DAPI and CTB-Alexa 647, resp. Confocal images were obtained with an LSM 880 point-scanning microscope with Airyscan (Zeiss) using a 20x (0.80 numerical aperture) or a 40x plan-apochromat objective (0.95 numerical aperture). Fluorophores were excited using a 405 nm diode, a 488 nm argon, a 561 nm diode-pumped solid-state (DPSS), and 633 nm helium-neon lasers. Detection intervals were set at (nm) 420-480 (DAPI), 500-550 (Alexa 488), 571-620 (mCherry), and 643-700 (Alexa 647).

Labelled cells were manually counted in 15 slices *per* animal, spanning the entire AP axis of the HIPP and 100 μm apart. Counts were then allocated to AP groups as dHIPP (−3.0 to −4.0 mm AP), diHIPP (−4.0 to −6.0 mm AP, above the rhinal fissure), and vHIPP (−4.0 to −6.0 mm AP, below the rhinal fissure). DAPI staining was used to identify hippocampal *strata*, and cell counts were grouped in CG, MCC, and RSC (cortical ‘region’), and, within each group, in HIPP AP groups (dHIPP, diHIPP, and vHIPP, ‘hippAP’) or according to *strata* (SO, SP, SR, SR-SLM, ‘strata’). Statistical significance of differences in CTB+ cell counts were tested using N-way ANOVA for either {‘region’, ‘hippAP’} or {‘region’, ‘strata’}. Bonferroni’s correction was applied in all *post-hoc* multiple comparisons throughout the manuscript.

For anterograde neuronal tracing, mCherry positive axons were manually counted using Fiji software on 11 slices *per* animal and per construct injected (CaMKIIa.hChR2 and CAG.hChR2). Of these slices (200 μm apart) 3 contained CG (subarea 24a), 4 MCC (subarea 24a’), and 4 (500 μm apart) RSC (subarea 29c). On each slice we counted the number of axons intersecting a grid of 8 lines orthogonal to L1 and 25 μm apart, drawn across a ROI of 255 μm x 1169 μm. On each individual slice, axon density in each layer (L1 to L6) was computed by dividing the total number of axons counted by its thickness (subareas 24a and 24a’: L1, 210 μm; L2, 65 μm; L3, 177 μm; L5, 210 μm; L6, 370 μm; subarea 29c: L1, 160 μm; L2, 45 μm; L3, 68 μm; L4, 68 μm; L5, 365 μm; L6, 273 μm). Densities were then grouped by MMC region and superficial (L1-L4) vs deep layer (L5-L6) (‘layer’) contingents. Differences in axon density were tested using N-way ANOVA {‘region’, ‘layer’}. Quantification of mCherry fluorescence along the cortical axis was computed as in (Morgenstern et al., 2016), with total values background corrected by subtraction of the mode, normalized, and averaged within cortical region across rats. Colocalization analysis was performed using the Colocalization Threshold plugin (Fiji) as in (Temido-Ferreira et al., 2018).

### Optogenetics and in vitro electrophysiology

Eight and 3 male Sprague-Dawley rats received two unilateral injections of either AAV_9_.CaMKIIa.hChR2-mCherry, or AAV_9_.CAG.hChR2-mCherry in diHIPP, resp (coordinates available in ST2), were decapitated 4 to 6 weeks post-injection, under deep isoflurane anesthesia, their brains quickly chilled for 3 min in ice-cold oxygenated (95% O_2_, 5% CO_2_) dissection solution [mmol/L: sucrose 110, KCl 2.5, CaCl_2_ 0.5, MgCl_2_ 7, NaHCO_3_ 25, NaH_2_PO_4_ 1.25, glucose 7 (pH 7.4)], trimmed to a 7 mm block, glued onto the stage of a vibratome (Leica VT1200S) using cyanoacrylate, and sliced in 400 μm thick coronal slices at an angle of 10° to the vertical. Slices were then transferred to a chamber containing oxygenated aCSF at 35°C, for 20-25 min, and then to a storage chamber containing oxygenated aCSF at RT (20-25°C) (based on Lu et al., 2014; Rombo et al., 2014, 2016).

After 45 min at RT slices were submerged inside a MEA2100-System^®^ (Multi Channel Systems) chamber laid over a 60-channel multi-electrode insert (60MEA200/30iR-Ti), perfused with 2 mL/min oxygenated aCSF at 30°C. Optogenetics-evoked fPSP were recorded using MC Rack software (Multi Channel Systems) under blue light stimulation triggered by a Grass Stimulus Generator TTL pulse driving an LED Plexon assembly. Remote expression of ChR2 and slice viability were assured by performing initial recordings on the hippocampal slice contralateral to the injected one, and electrical stimulation, resp. Optimal stimulation sites were determined by placing the fiberoptic over different regions of the slice until maximal network responses, driven at 30 mW of total LED power, 100ms pulse length and a 10s IPI. After stable responses for 10 min the experimental protocol was started. CaMKII slices were sequentially perfused with aCSF (condition CTCAM), and the following additive drug treatments, each for 20 min: T1_CAM_ (PTX, 50 μM), T2_CAM_ (the previous plus CNQX, 20 μM), T3_CAM_ (idem, APV, 50 μM). Likewise CAG slices were sequentially recorded in aCSF (condition CT_CAG_), T1_CAG_ (CNQX), T2_CAG_ (plus APV) T3_CAG_ (plus PTX), to isolate remote GABAa inputs, as explained in the main text. Responses were amplified and digitized at 50 kHz, and the slice position in the MEA was captured using a monochromatic camera (Celestron). The expression of the viral construct was verified histologically for each slice. This protocol was based on (Hass and Glickfeld, 2016; Lu et al., 2014).

### Data analysis and quantification

Downsampled (2 kHz) responses were baseline-corrected, and further analyses were done using code written in MATLAB. For each channel we compared the slopes in the last 2 min of CT_CAM_ with the last 2 min of T3_CAM_, within channels. Slopes were computed as the voltage differences between t0 (stim onset) and t1 (+60ms) divided by time (Figure 4D). Wilcoxon rank sum test was used to determine statistical significance of pre vs post treatment responses. For these channels dimmed to have a synaptic response, we computed the slope and amplitude (max difference between t0 and t0+160ms) of the response difference. Synaptic response slopes and amplitudes were grouped by MMC region (‘region’) and superficial vs deep (‘layer’), for N-way ANOVA {‘region’, ‘layer’}. Responses in CAG slices were analyzed as above, with modifications. Channels with significant responses were selected by comparing the slopes in CT_CAG_ *versus* T2_CAG_. Excitatory responses were obtained by subtracting T2_CAG_ from CT_CAG_, inhibitory responses by subtracting T3_CAG_ from T2_CAG_. Synaptic responses from RSC were grouped in superficial and deep responses (‘layer’), and, within each group, in excitatory and inhibitory (‘type’) for N-way ANOVA {‘layer’, ‘type’}.

### In vivo electrophysiology

Animals were implanted with a hyperdrive with 32 independently movable tetrodes (Liang et al., 2017). MMC was targeted with a linear array of 19 tetrodes (TT) implanted at +2.0 mm A/P and +0.5 mm M/L, CA1 with a 7-11 TT array −3.0 mm A/P and +2.0 mm M/L. Surgery was performed as above with some modifications as in (Remondes and Wilson, 2015). The hyperdrive was lowered with TT sticking out ventrally (1.5 mm for MMC and 2.0 mm for CA1). The hyperdrive was secured to the screws and bone with dental cement and the surgical wound was closed. Rats were allowed to recover for one week post-surgery.

Data included in this manuscript was acquired from awake rats in various behavioral states, namely running a delayed non-matching to trajectory (DNMT) task as in (Yamamoto and Tonegawa, 2017), open field exploration sessions with and without reward, and free exploration of a T-maze apparatus. Continuous LFP signals were recorded at 30kHz using the Intan’s RHD2164 boards, and Open Ephys (Siegle et al., 2017). XYT position was acquired at 30 fps using a Flea 3 Point Grey camera tracking an LED placed on the hyperdrive. Bonsai Software run in the acquisition computer managed all data acquisition and writing processes (Lopes et al., 2015). Raw data was band-pass filtered between 700 Hz and 8 kHz for spike waveform extraction, and between 0.1 and 8 kHz for LFP analysis. Action potentials were assigned to individual cells by offline clustering based on spike amplitudes, using UltraMegaSort 2000 (Hill et al., 2011). Subsequent analyses employed functions from the Chronux toolbox and code written in Matlab (Mathworks, Natick, MA).

Spikes from each brain region were binned at 10ms into a MUA vector per region and dataset. Analyzed data epochs corresponded to +/− 1 sec of enhanced HIPP spiking activity, defined as bins with over 4 spikes. Cortical binned MUA response, trigger-point response, spectra and coherence with HIPP MUA were computed, the latter two frequencies to a maximum of 50Hz (given MUA bin size of 10ms), in 50ms-stepping windows of 500ms duration. Such spectrograms and coherograms were normalized by a pre-trigger 0.5 seconds baseline, and pre vs post HIPP-trigger magnitudes were compared across relevant frequency bands, and MMC regions using N-way ANOVA with the appropriate factors. Presence of tetrodes was verified using histology, records of daily adjustment, and visually during surgery.

## Author contributions

EF-F and MR designed the experiments and analyzed the data, EF-F, CQ and MR conducted the experiments and MR wrote the manuscript.

## Acknowledgements

This work was supported by Fundação para a Ciência e Tecnologia – Portugal with a PhD fellowship to EFF, an Exploratory Grant and an IMM Director's Fund Award to MR, who holds an Investigator FCT Position at IMM. Marta Moita, Joao Peca, and Rui Costa provided us with optogenetic tools used in this study, Leopoldo Petreanu guidance in viral injection procedures, Ana Sebastiao and Luisa Lopes assisted us with *in vitro* electrophysiology setups, IMM's Bioimaging, Hystology, and Animal facilities for help and assistance, and to all the members of the Remondes Lab for fruitful discussions.

