## Supplementary Figure 1 for "A Gradient of Hippocampal Inputs to the Medial Mesocortex"

### Supplementary Figure 1
#### *(referred to figure 3)*



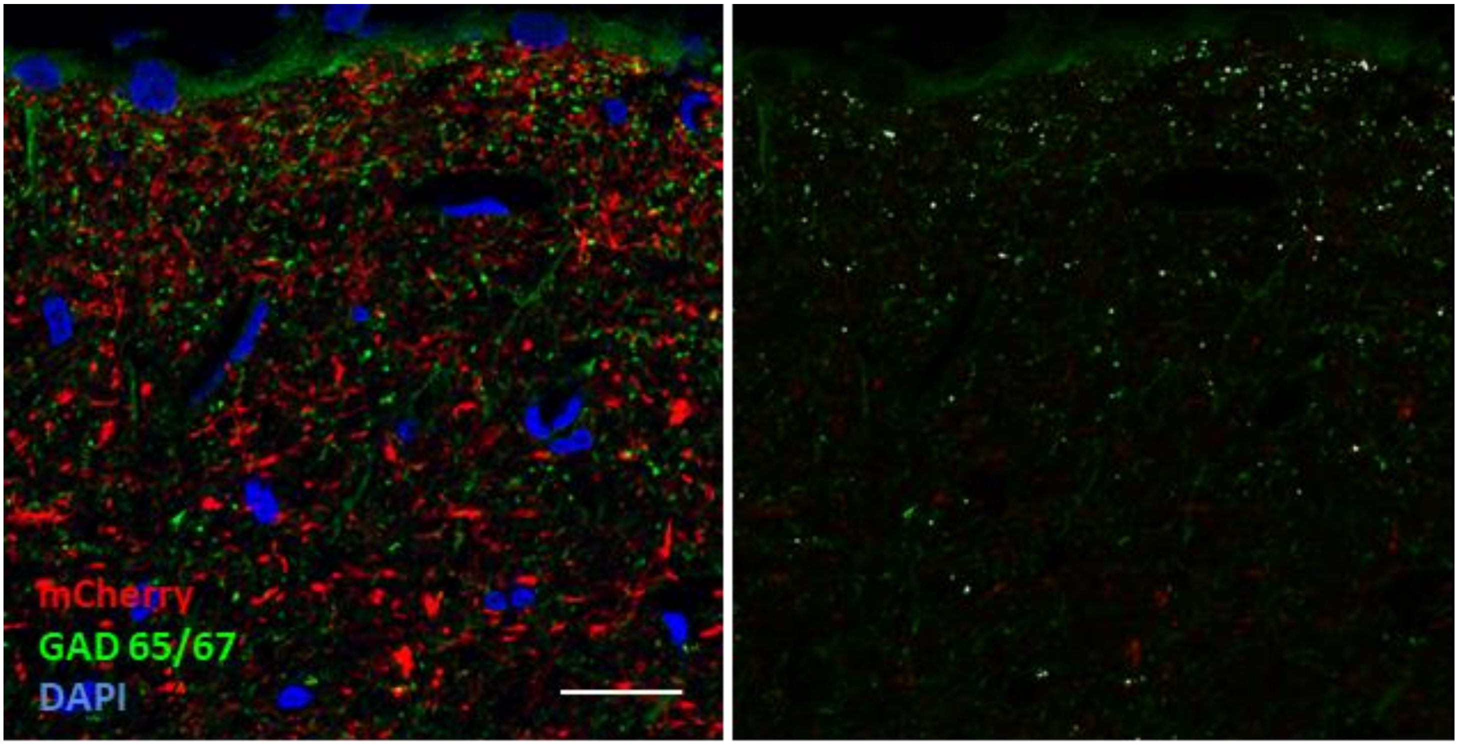


Supplementary Figure 1 – GAD+ puncta in RSC significantly colocalize with labelled Hippocampal axon terminals, strongly suggesting that these are of inhibitory terminals (please see Supp Table 1 for quantifications). Single plane confocal picture from one example rat showing mCherry+ hippocampal axons and GAD65/67+ puncta in layer 1 of RSC (left). Co-occurring objects are identified as gray dots on the right panel. Scale bar: 20 µm.
