## Supplementary Figure 2 for "A Gradient of Hippocampal Inputs to the Medial Mesocortex"

### Supplementary Figure 2
#### *(referred to figure 6)*

***Ferreira-Fernandes et al 2018***

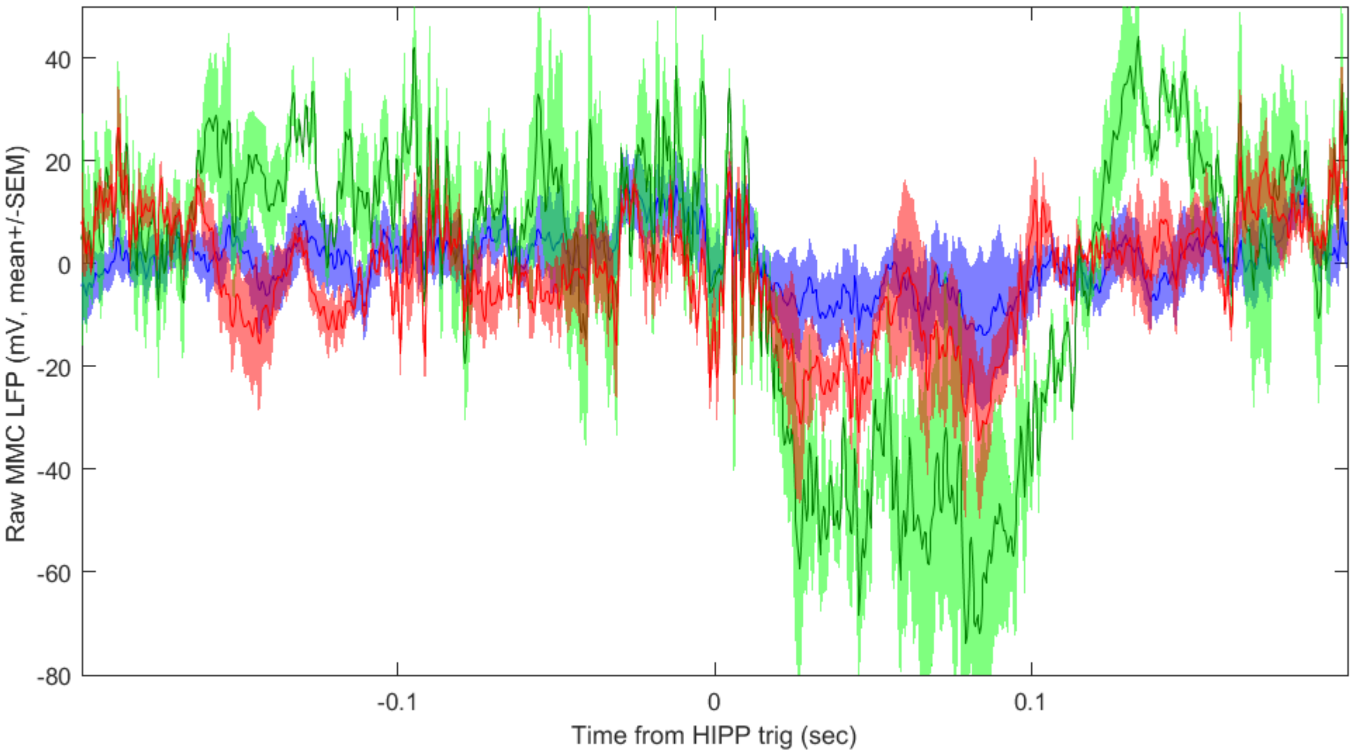


Supplementary Figure 2 – Raw LFP from the three MMC regions at the same epochs triggered by HIPP spikes defined previously, color coded as before. Note the presence of complex deflections triggered by increases in HIPP MUA, of somewhat lower amplitude in RSC.
