## Supplementary Figure 3 for "A Gradient of Hippocampal Inputs to the Medial Mesocortex"

**Supplementary Figure 3**
***(referred to Figure 8)***



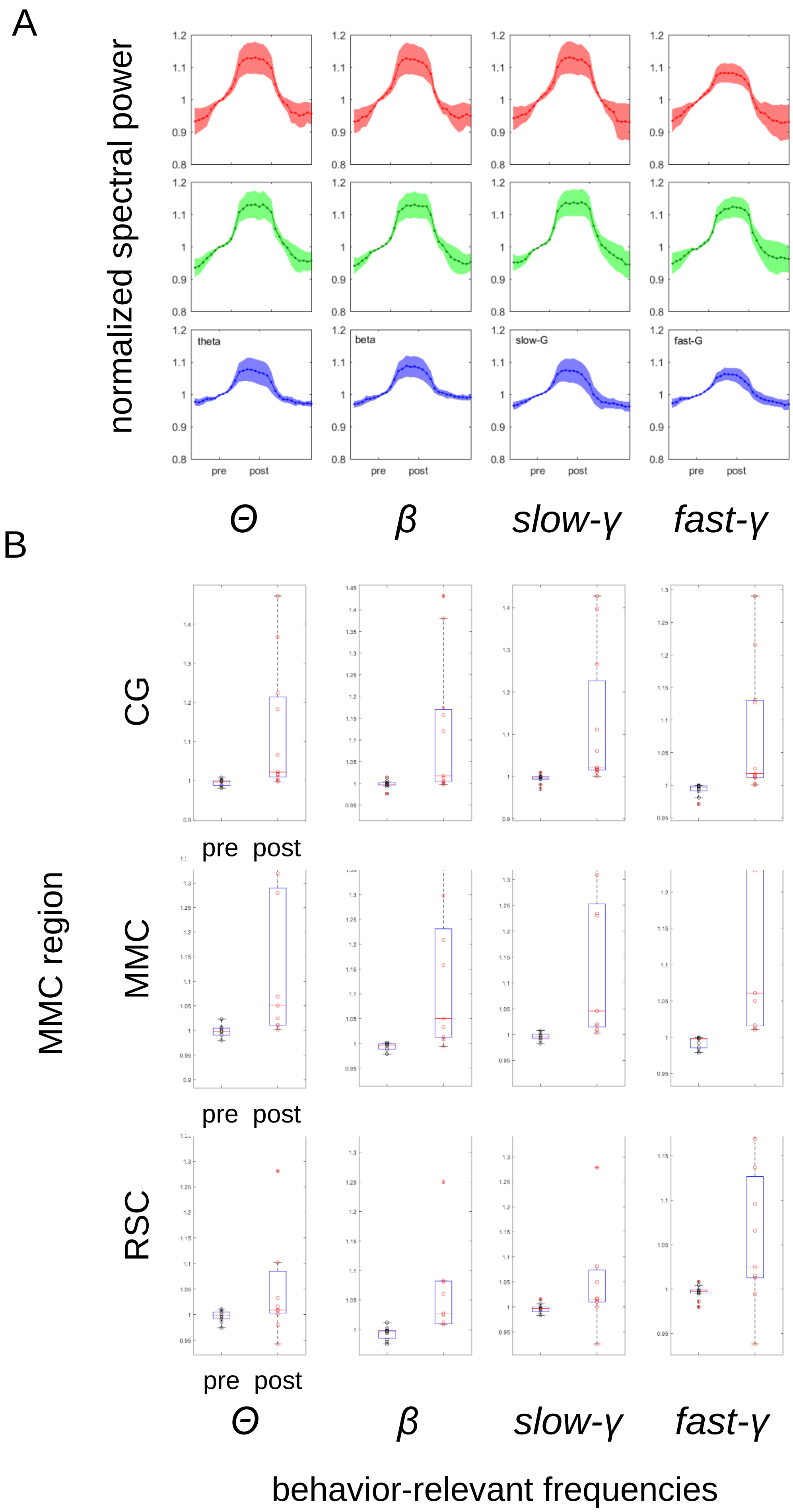

**Supplementary Figure 3 -** Medial mesocortical responses to HIPP spikes at behavior-relevant frequencies (A) Color-coded plots depict the quantification of HIPP-triggered MMC power at relevant frequency bands, normalized to a pre-trigger baseline of 0.5 seconds. (B) There is a significant HIPP-triggered power increase at all frequencies regardless of cortical region analyzed (n-way ANOVA with factors pre- vs post- HIPP trigger, cortical region, and frequency, $F(1,224)=78.06$, $n=11$ datasets, we present here boxplots for each condition for informative purposes).
