## Supplementary Table 1 for "A Gradient of Hippocampal Inputs to the Medial Mesocortex"

Supplementary Table 1 – Absolute numbers of CTB positive neurons grouped by MMC injection site, hippocampal anatomical level (top) and hippocampal stratum (*so, sp, sr, sr/slm*)

**CTB POSITIVE NEURONS**

**AVERAGE ± STANDARD DEVIATION**

**Injection Sites**

dHIPP

diHIPP

vHIPP

**CG**

**(A, B1, B2, B3; n = 4)**

(

2, 8, 3, 0

)

3.25 ± 3.40

(

155, 63, 175, 288)

170.25 ± 92.42

(76, 21, 79, 65)

60.25 ± 26.85

**MCC**

**(C1, C2, C3; n = 3)**

(

47, 1, 3

)

17 ± 26

(154, 49, 49)

84 ± 60.62

(33, 14, 8)

18.33 ± 13.05

**RSC**

**(D, E, F; n = 3)**

(

32, 46, 41

)

39.67

± 7.09

(139, 227, 59)

141.67 ± 84.03

(48, 0, 0)

16 ± 27.71

**CTB POSITIVE NEURONS**

**AVERAGE ± STANDARD**

**DEVIATION**

**Injection Sites**

SO

SP

SR

SR/SLM

**CG (A, B1, B2, B3; n = 4)**

(2, 2, 1, 6)

2.75 ± 2.22

(227, 65, 252, 317)

215.25 ± 107.11

(4, 6, 4, 27)

10.25 ± 11.21

(0, 19, 0, 3)

5.5 ± 9.11

**MCC (C1, C2, C3; n = 3)**

(3, 0, 1)

1.33 ± 1.53

(137, 63, 58)

86 ± 44.24

(25, 1, 1)

9 ± 13.86

(69, 0, 0)

23 ± 39.84

**RSC (D, E, F; n = 3)**

(5, 21, 1)

9 ± 10.58

(108, 11, 95)

71.33 ± 52.65

(36, 53, 2)

30.33 ± 25.97

(70, 188, 2)

86.67 ± 94.11

**Supplementary Table 1	*Ferreira-Fernandes et al 2018***

***(referred to figure 1)***
