## Supplementary Table 2 for "A Gradient of Hippocampal Inputs to the Medial Mesocortex"

Supplemental Table 2 – Colocalization analysis quantification. tM thresholded Mander’s split colocalization Coefficient; Th, Costes’ threshold; r<th correlation below the Costes’ threshold. All values are AVG+/-SD.

**Supplemental Table 2	*Ferreira-Fernandes et al 2018***

***(referred to figure 3)***

**Layers**

**tM**

**GAD**

**T**

**h**

**mCherry**

**Th**

**GAD**

**r<th**

**1 (n = 3)**

0.47 ± 0.17

6.44 ± 2.51

41.11 ± 40.16

0.005 ± 0.016

**3/4 (n = 3)**

0.10 ± 0.08

92.44 ± 121.93

8.44 ± 2.83

0.001 ± 0.006

**5 (n = 3)**

~0

<0

<0

-
