## Supplementary figures and images for "A Gradient of Hippocampal Inputs to the Medial Mesocortex"

### Supplementary Table 3

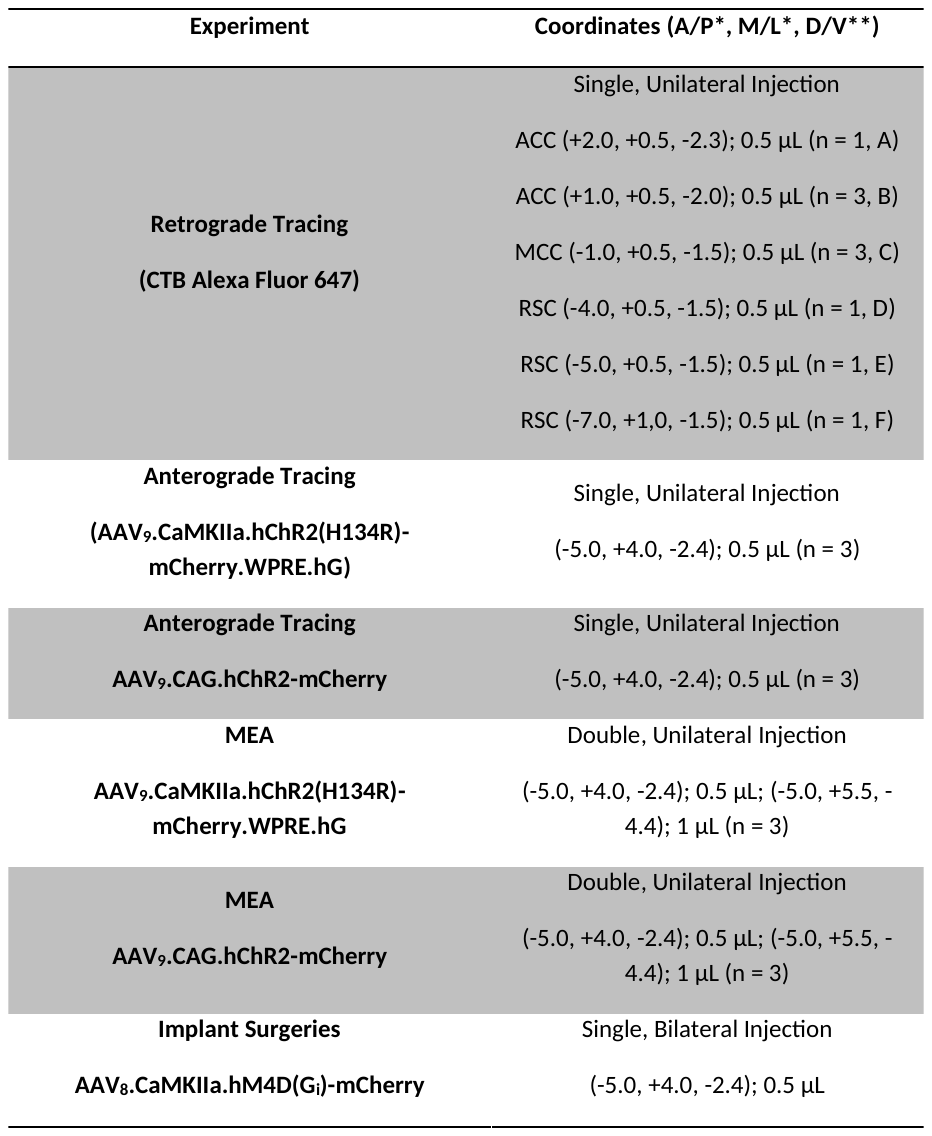

**Supplemental Table 3	*Ferreira-Fernandes et al 2018***

***(referred to Methods)***
